# Bio-informatic Analysis of Missense Single Nucleotide Polymorphisms (SNPs) in Human CD38 Gene Associated with B-Chronic Lymphocytic Leukemia

**DOI:** 10.1101/2020.08.12.241976

**Authors:** Hiba Awadelkareem Osman Fadl, Abdelrahman Hamza Abdelmoneim, Sahar Gamal Elbager

## Abstract

**Background:** CLL: Chronic lymphocytic leukemia is a chronic type of haematological malignancies that evoked from lymph proliferative origin of bone marrow and secondary lymphoid tissue, resultant in proliferation and progressive accumulation of distinct monoclonal CD5 /CD19 /CD23 B lymphocytes in the bone marrow, peripheral blood, and lymphatic organs. **CD38** is a multifunctional ecto-enzyme, known to be a direct contributor in pathogenesis of CLL by poorly understood mechanism. Even though, it highly expressed in CLL. At specific position of CD38 gene sequence, substitution of single nucleotide may result in change in amino acid that ends by consequent alteration of protein structure.

**Aim:** To study CD38 polymorphism and to predict its effect on structure and subsequently function of CD38 molecule.

**Methodology and Result:** The bioinformatic analysis of CD38 gene had been carried out by using several soft wares. Functional analysis by SIFT, Polyphen2, and PROVEAN reveled 12 deleterious SNPs. These SNPs were further analyzed by SNAP2, SNP@GO. PMut, STRING and other soft wars. Furthermore, Stability analysis was done using I-Mutant and MUpro software where seven SNPs were found to decrease the stability of the protein by I-Mutant, while two SNPs increase it. At the same time, eight SNPs were found to decrease the stability by Mupro software while only one SNP is predicted to increase it. Finally, Physiochemical analysis was done using Project Hope.

**Conclusion:** In summary, CD38 genotype seems to have twelve SNP that possibly will result in deleterious effect on Protein Structure. This genetic variation eventually will lead to alteration in potential molecule functions. Which effect the progression of CLL By the end.

## Introduction

Chronic lymphocytic leukemia (CLL) is a chronic type of hematological malignancies that evoked from lymph proliferative origin and derived from bone marrow and secondary lymphoid tissue which accompanied by exaggerated monoclonal proliferation of mature B lymphocytes**^(1)^**, result in progressive accumulation of distinct monoclonal CD5+/CD19+/CD23+ B lymphocytes in the bone marrow, peripheral blood, and lymphatic organs**^(2)^**. On an other hand, CLL is the most common type of leukemia among adults**^(3)^**. furthermore, It is a Malignancy that mostly affects elders; 80% of those patients are aged more than 50 years **^(4,5)^** although, B-CLL is molecularly heterogeneous disease^(6)^ but the exact etiology of this leukemia is not well understood.**^(7)^** In fact, The clinical course for B-CLL is highly variable, fortunately this matter recently were solved by the analysis of novel molecular prognostic factors including CD38 and ZAP-70 expression.**^(8)^**.

CD38(Cluster Of Differentiation 38) also known as ADPRC1, it is a type II trans membrane glycoprotein in nature which is defined as a multifunctional ecto-enzyme that expressed normally at high levels on plasma and nuclear membranes of B cell precursors, germinal center B cells, and plasma cells, with low expression on circulating B cells.The gene that encodes human CD38 has been mapped in the short arm of chromosome 4 (4p15) with its coding sequence organized in eight exons and more than 98% of the gene is represented by introns **^(9).^**

### CD 38 Structure

As a Trans membrane protein, full-length human CD38 comprised from 300 amino acid residues connecting to three domains: a short intracellular domain, a single trans membrane helix domain, and a large extracellular domain. The distribution of amino acids among domains as following: Cytoplasmic domain from 1-21, Trans membrane domain 22-42 and extracellular domain 43-300 amino acid.

As known, intracellular domain act as enzyme and extra cellular domain as receptor. The extracellular domain is an ecto-enzyme (EC3.2.2.5) which produce cyclic ADP-ribose (cADPR) and nicotinic acid adenine dinucleotide phosphate (NAADP) from its substrates NAD and NADP, in that order. **^(10)^**

The biological functions of CD38 molecule is varied depends on the cells type and CD38 maturation state. CD38 is known to be functionally active in signal transduction, cell adhesion and calcium signaling. The binding to the ligand CD31, initiates a signaling cascade that includes phosphorylation of sequential intracellular targets and increases cytoplasmic Ca**^+2^** levels, mediating different biological CD38 (CD38 molecule)**^(11).^** Additionally, another events of CD38 involvement in biological activities had been noted (e.g., activation, proliferation, apoptosis, cytokines secretion and homing), which depends on the cells type. CD38 accomplished its role As an enzyme, generate ng cADPR, ADP-ribose and NAADP CD38 as a products of AD+/NADP+ metabolism. These products bind different receptors and channels (IP3 receptors IP3R, Ryanodine receptor RyR and Transient receptor potential cation channel subfamily M member 2 TRPM2) and are involved in the regulation of intracellular Ca**^+2^** and activation of critical signaling pathways connected to the control of cell metabolism, genomic stability, apoptosis, cell signaling, inflammatory response and stress tolerance.**^(12)^** Several studies reported over expression of CD38 in CLL.**^(13)^** Recent findings have suggested that CD38 is not only a prognostic marker, but also the key component of the pathogenic network underlying B CLL and T. **^(14)^**

Single nucleotide polymorphisms (SNPs) in CD38 gene were detected in numerous haematological disorders such as CLL, multiple myeloma, Correlectal cancer and SLE as well as other disorders like Diabetes and AutismREF. Most of the studies were used molecular tools, till now no studies talking a bioinformatics analysis of CD38. In the current study we performed a comprehensive bioinformatics analysis to predict the possible SNPs in CD38 gene and assessing its effects on the structure and function of **ADPRC1** protein.**^(15–19)^**

## Method

### 2.1. Retrieving nsSNPs

SNPs associated with CD38 gene were obtained from the Single Nucleotide Polymorphism database (dbSNP) in the National Center for Biotechnology Information (NCBI). (http://www.ncbi.nlm.nih.gov/snp/).

The sequence and natural variants of CD38 protein were obtained from the UniProt database as it considered as the most reliable and unambiguous database for protein sequences.**^(20)^**(https://www.uniprot.org/).

### 2.2. Identifying the most damaging nsSNPs and disease related mutations

Five different bioinformatics tools were utilized to predict functional effects of nsSNPs obtained from dbSNP database on the protein. These algorithmic include: SIFT, PROVEAN, PolyPhen-2, SNAP2 and PhD-SNP.

The SNPs predicted deleterious by at least three softwares were considered high risk nsSNPs and investigated further.

#### 2.2.1. SIFT Server

Phenotypic effects of amino acid substitution on protein function were predicted by using Sorting Intolerant From Tolerant (SIFT) server, which is a powerful tool used to fulfill this purpose. A list of nsSNPs (rsIDs) from NCBI's dbSNP database was submitted as a query (original) sequence to SIFT to predict tolerated and deleterious substitutions for every position of a protein sequence. . (Available at: http://sift.bii.a-star.edu.sg/). **^(21–24)^**

#### 2.2.2. Provean Server

(Protein Variation Effect Analyzer) is the second software tool used. It also predicts the effect of an amino acid substitution on the biological function of a protein. It predicts the damaging effects of any type of protein sequence variations to not only single amino acid substitutions but also in-frame insertions, deletions, and multiple amino acid substitutions. (Available at: http://provean.jcvi.org/index.php). **^(25–26)^**

#### 2.2.3. Polyphen-2 Server

Polymorphism Phenotyping v2.0 (PolyPhen-2) is another online tool that predicts the possible effects of an amino acid substitution on the structure and function of the protein. The results are classified into “PROBABLY DAMAGING” that is the most disease causing with a score near to 1 (0.7-1), “POSSIBLY DAMAGING” with a less disease causing ability with a score of 0.5-0.8 and “BENIGN” which does not alter protein functions with a score closer to zero; (0-0.4). (Available at: http://genetics.bwh.harvard.edu/pph2/). **^(27–29)^**

#### 2.2.4. SNAP2

It is a trained functional analysis web-based tool that differentiates between effect and neutral SNPs by taking a variety of features into account. SNAP2 has an accuracy (effect/neutral) of 83%. It is considered an important and substantial enhancement over other methods.**^(30)^**

#### 2.2.5. CONDEL

Condel is a method to assess the outcome of non-synonymous single nucleotide variants SNVs using a consensus deleteriousness score that combines various tools (SIFT, Polyphen2, <u>MutationAssessor</u>, <u>and FATHMM</u>). The result of the prediction are either deleterious or neutral with a score ranging from zero to one.**^(31)^**

### 2.3 Prediction of Disease Associated SNPs

#### 2.3.1. SNPs & Go server

An online web server that used to ensure the disease relationship with the studied single nucleotide polymorphisms SNPs. It gives three different results based on three different analytical algorithms; Panther result, PHD-SNP result, SNPs &GO result. (Available at: http://snps-and-go.biocomp.unibo.it/snps-and-go/).**^(32)^**

#### 2.3.2. PMUT Server

PMUT is a powerful web-based tool used for the prediction of pathological variants on proteins. The prediction results are catogrized as “Neutral” or “Disease”. It is available at (http://mmb.irbbarcelona.org/PMut).**^(33)^**

### 2.4 Prediction of nsSNPs Impact on the Protein Stability by I-Mutant2.0

#### 2.4.1 I-Mutant

A online web-tool that is used for the prediction of protein stability changes upon single point mutations, determining whether the mutation increases or decreases the protein stability. (Available at: http://gpcr.biocomp.unibo.it/cgi/predictors/I-Mutant2.0/I-Mutant2.0.cgi). **^(34)^**

#### 2.4.2 Mupro

It is a support vector machine-based tool for the prediction of protein stability changes upon nonsynonymous SNPs. +e value of the energy change is predicted, and a confidence score between −1 and 1 for measuring the confidence of the prediction is calculated. A score <0 means the variant decreases the protein stability; conversely, a score >0 means the variant increases the protein stability. **^(35)^**

### 2.5 Identification of Functional SNPs in Conserved Regions by ConSurf

#### 2.5.1 ConSurf analysis

It is a web server that offers evolutionary conservation summaries for proteins of known structure in the protein data bank. ConSurf spots the parallel amino acid sequences and runs multialignment methods.**^(36)^**

### 2.6 Analyzing the Effect of nsSNPs on Physiochemical Properties by Mutpred

#### 2.6.1 MutPred server

MutPred2 is a standalone and web application developed to classify amino acid substitutions as pathogenic or benign in human. In addition, it predicts their impact on over 50 different protein properties and, thus, enables the inference of molecular mechanisms of pathogenicity. **^(37)^**

### 2.7 Prediction of structural effect of point mutation on the protein sequence using Project HOPE

#### Project HOPE

It is an online web-server used to analyze the structural and functional variations on protein sequence that have been resulted from single amino acid substitution. It searches protein 3D structures by collecting structural information from a series of sources, including calculations on the 3D coordinates of the protein and sequence annotations from the UniProt database. Protein sequences are submitted to project HOPE server then HOPE builds a complete report with text, figures, and animations. It is available at (http://www.cmbi.ru.nl/hope).**^(38)^**

### 2.8 Protein-protein interactions analysis

The **STRING** database aims to collect, score and integrate all publicly available sources of protein– protein interaction information, and to complement these with computational predictions.

#### KEGG

(Kyoto Encyclopedia of Genes and Genomes: http://www.genome.jp/kegg/)**^(39)^** is a database resource which embraces genomic, chemical and systemic functional information, gene lists from fully sequenced genomes that interrelated to higher-level systemic functions of the cell, the organism and the ecosystem. KEGG is used broadly as a reference knowledge base designed for explanation of large-datasets that created by genome sequencing and other experimental technologies to support basic research, and practical applications of human diseases, drugs and other health-related matters.

## 3. Result

### 3.1 SNP Dataset from dbSNP

The reference sequence of CD38 protein (ID: P28907) was retrieved from National center NCBI databases along with related single nucleotide polymorphisms SNPs.

### 3.2. Prediction of Functional Mutations

Furthermore these SNPs(51) were submitted to SIFT server for prediction of function effect where 12 SNP were found to have deleterious effect on Protein (table1). Further functional analysis was done using polyhen2, PROVEAN and SNAP2, where 9 SNPs were found deleterious by the four previous software (table 2).

**Table 1:** functional analysis of SNPs in CD38 protein using SIFT, POLYPHEN2, PROVEAN, SNAP2, and CONDEL.

| SNP | AMINO ACID | SIFT_SCORE | SIFT_PREDICTION | POLYPHEN2_SCORE | POLYPHEN2_PREDICTION | PROVEAN_score | PROVEANPrediction | SNAP2_SCORE | SNAP2 | CONDEL | CONDEL_LABEL |
| --- | --- | --- | --- | --- | --- | --- | --- | --- | --- | --- | --- |
| rs1800561 | R140W | 0.012 | DELETARIOUS | 0.999 | probably damaging | -4.403 | Deleterious | 93 | effect | 0.409454 | Neutral |
| rs138866692 | T148M | 0.006 | DELETARIOUS | 1 | probably damaging | -3.866 | Deleterious | 67 | effect | 0.55141 | DELETARIOUS |
| rs149299047 | K234N | 0.012 | DELETARIOUS | 0.998 | probably damaging | -3.692 | Deleterious | 71 | effect | 0.553697 | DELETARIOUS |
| rs150171847 | S193C | 0 | DELETARIOUS | 1 | probably damaging | -4.771 | Deleterious | 72 | effect | 0.516534 | Neutral |
| rs199998792 | D147G | 0.018 | DELETARIOUS | 0.999 | probably damaging | -5.34 | Deleterious | 87 | effect | 0.4864 | Neutral |
| rs201683843 | F284C | 0.008 | DELETARIOUS | 1 | probably damaging | -3.036 | Deleterious | 10 | effect | 0.419877 | Neutral |
| rs201816957 | D179G | 0.035 | DELETARIOUS | 0.999 | probably damaging | -3.79 | Deleterious | 38 | effect | 0.377277 | Neutral |
| rs369214160 | T144N | 0.014 | DELETARIOUS | 1 | probably damaging | -3.596 | Deleterious | 50 | effect | 0.581067 | DELETARIOUS |
| rs369674233 | P98L | 0.007 | DELETARIOUS | 1 | probably damaging | -9.568 | Deleterious | 48 | effect | 0.502413 | Neutral |

**Table(2):** functional analysis of SNPs in CD38 protein using : SNP@GO, PHD-SNP, PANTHER, and PMut.

| AMINO_<br>ACID | Predicti<br>on<br>PANT<br>HER | R<br>I | Probab<br>ility | Predict<br>ion<br>SNP &<br>GO | R<br>I | Probab<br>ility | Predict<br>ion-PHD-SNP | R<br>I | Probab<br>ility | Pmut<br>predict<br>ion | Pmu<br>t<br>resu<br>lt |
| --- | --- | --- | --- | --- | --- | --- | --- | --- | --- | --- | --- |
| R140W | Disease | 8 | 0.925 | Neutra<br>l | 5 | 0.237 | Neutral | 4 | 0.288 | Neut<br>ral | **0.08**<br>(96<br>%) |
| T148M | Disease | 1 | 0.567 | Neutra<br>l | 6 | 0.224 | Disease | 0 | 0.504 | Neut<br>ral | **0.13**<br>(95<br>%) |
| K234N | Disease | 3 | 0.674 | Neutra<br>l | 1 | 0.461 | Disease | 1 | 0.544 | Neut<br>ral | **0.22**<br>(92<br>%) |
| S193C | Disease | 7 | 0.844 | Diseas<br>e | 2 | 0.613 | Disease | 4 | 0.708 | Dise<br>ase | **0.85**<br>(91<br>%) |
| D147G | Disease | 5 | 0.746 | Diseas<br>e | 2 | 0.622 | Disease | 4 | 0.685 | Dise<br>ase | **0.53**<br>(80<br>%) |
| F284C | Disease | 7 | 0.839 | Neutra<br>l | 2 | 0.409 | Disease | 1 | 0.557 | Neut<br>ral | **0.16**<br>(93<br>%) |
| D179G | Disease | 2 | 0.579 | Neutra<br>l | 2 | 0.375 | Disease | 0 | 0.51 | Neut<br>ral | **0.23**<br>(91<br>%) |
| T144N | Disease | 1 | 0.534 | Neutra<br>l | 2 | 0.38 | Disease | 2 | 0.586 | Neut<br>ral | **0.31**<br>(89<br>%) |
| P98L | Disease | 8 | 0.879 | Diseas<br>e | 2 | 0.579 | Disease | 2 | 0.58 | Dise<br>ase | **0.63**<br>(84<br>%) |

### 3.3 Prediction of Disease Associated SNPs

### 3. 4 Prediction of nsSNPs Impact on the Protein Stability

Stability analysis was done using Imutant and MUpro software where seven SNPs were found to decrease the stability of the protein by Imutant, while two SNPs(T148M, T144N) increase it. At the same time, eight SNPs were found to decrease the stability by Mupro software while only one SNP(P98L) are predicted to increase it.(Table3).

**Table3.** Analysis of ADPRC 1 using I-Mutant Suite.

| Residue | Imutant<br>Prediction Effect | Value | Mupro<br>Prediction Effect<br>(Stability) | Value |
| --- | --- | --- | --- | --- |
| R140W | Decrease | 0.13 | Decrease | -0.743 |
| T148M | Increase | -0.07 | Decrease | -0.471 |
| K234N | Decrease | -0.23 | Decrease | -0.666 |
| S193C | Decrease | -0.60 | Decrease | -0.168 |
| D147G | Decrease | -0.92 | Decrease | -1.514 |
| F284C | Decrease | -1.90 | Decrease | -1.285 |
| D179G | Decrease | -1.39 | Decrease | -1.131 |
| T144N | Increase | -0.52 | Decrease | -0.770 |
| P98L | Decrease | -0.30 | Increase | 0.217 |

### 3. 5 Analysis of Conservation profile

### 3.6 Analyzing the Effect of nsSNPs on Physiochemical Properties Mutpred

Physiochemical analysis was done using Project Hope. A summary of the predicted effect on the structure and physiochemical priorities were summarized in (Table 4).

**Table 4.**
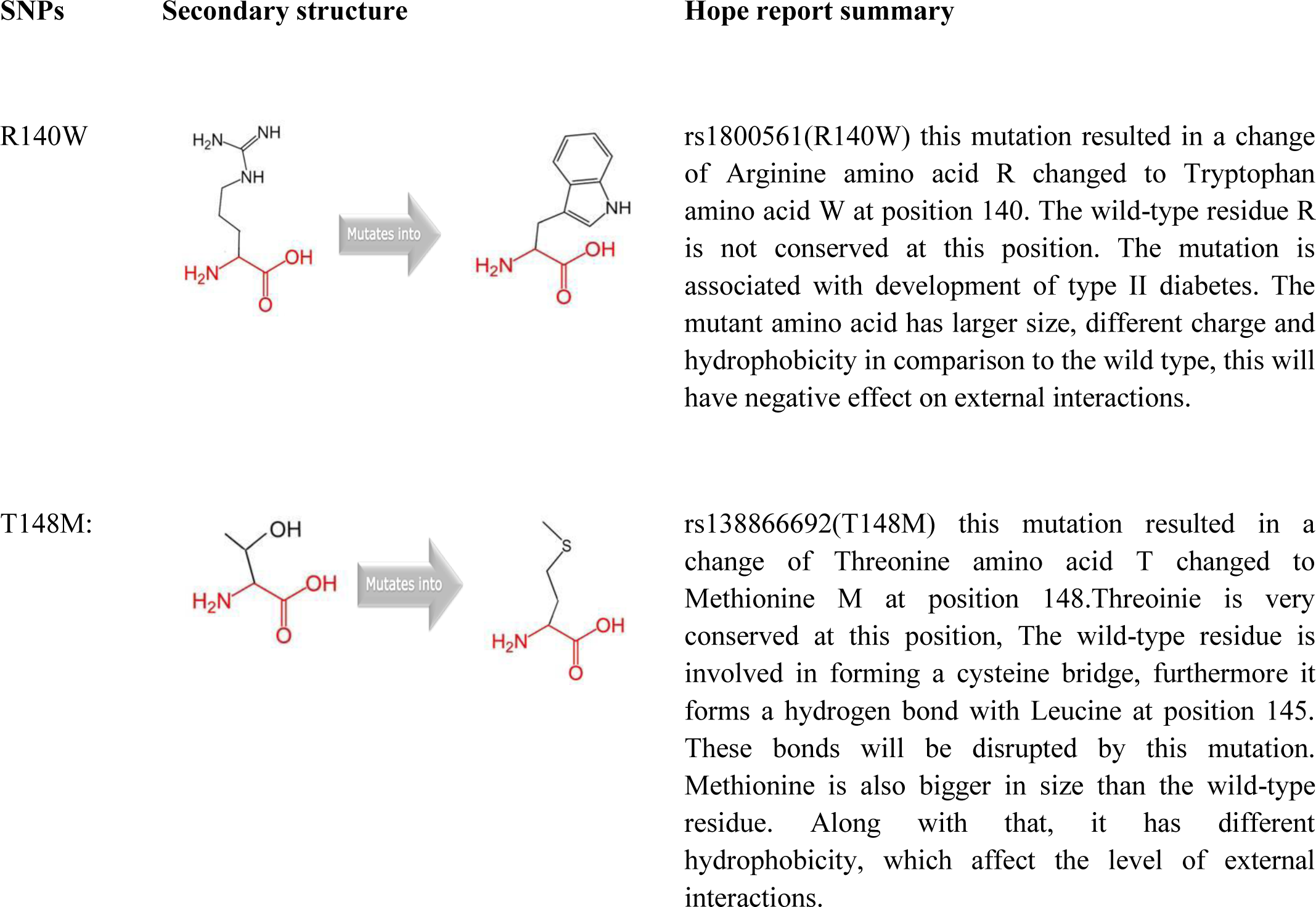
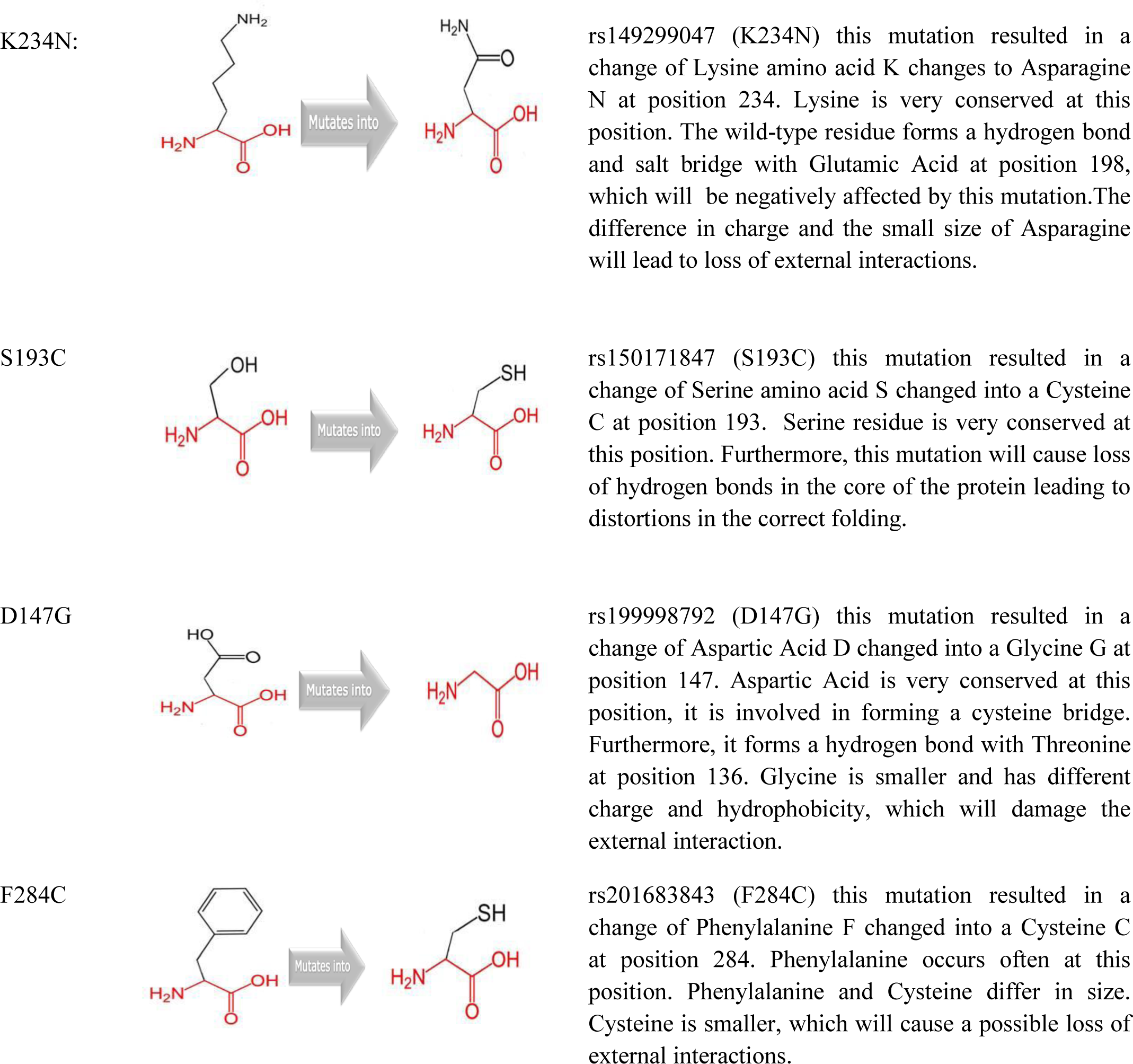
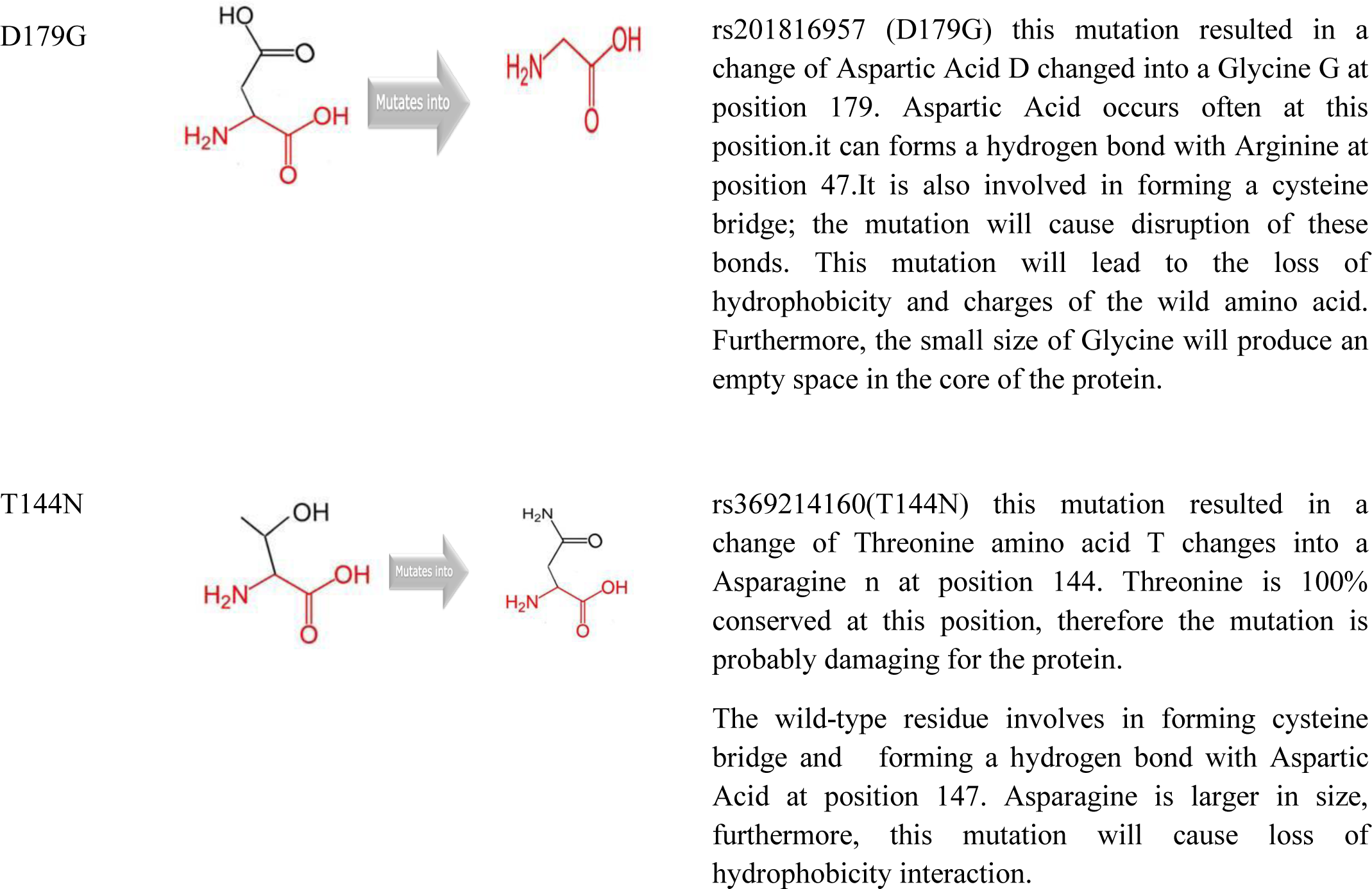
the physiochemical proprieties induced by the SNPs in CD38 gene as predicted by Hope software.

#### Analysis of Conservation profile

ConSurf analysis predicted P98L, T144N and D147G to be exposed highly conserved and exposed, i.e., a functional residue, which suggests that these positions are important for the ADPRC 1 function. ConSurf analysis also predicted T148M and S193C to be buried and conserved residue, i.e., a structural residue (Fig.1, Table5).

**Fig1..**
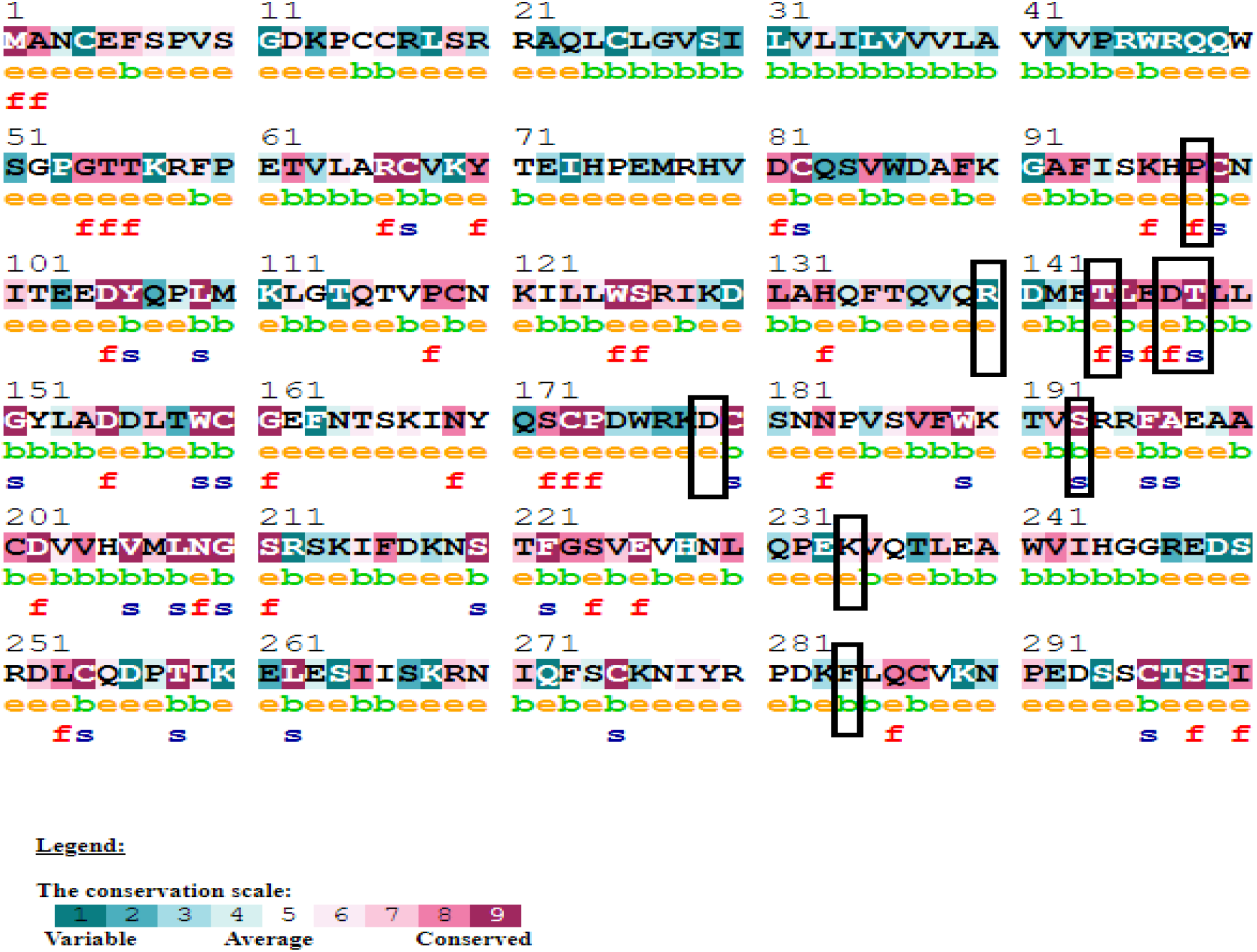
Analysis of evolutionary conserved amino acid residues of ADPRC 1 by ConSurf. The color coding bar shows the coloring scheme representation of conservation score.

**Table 5:**
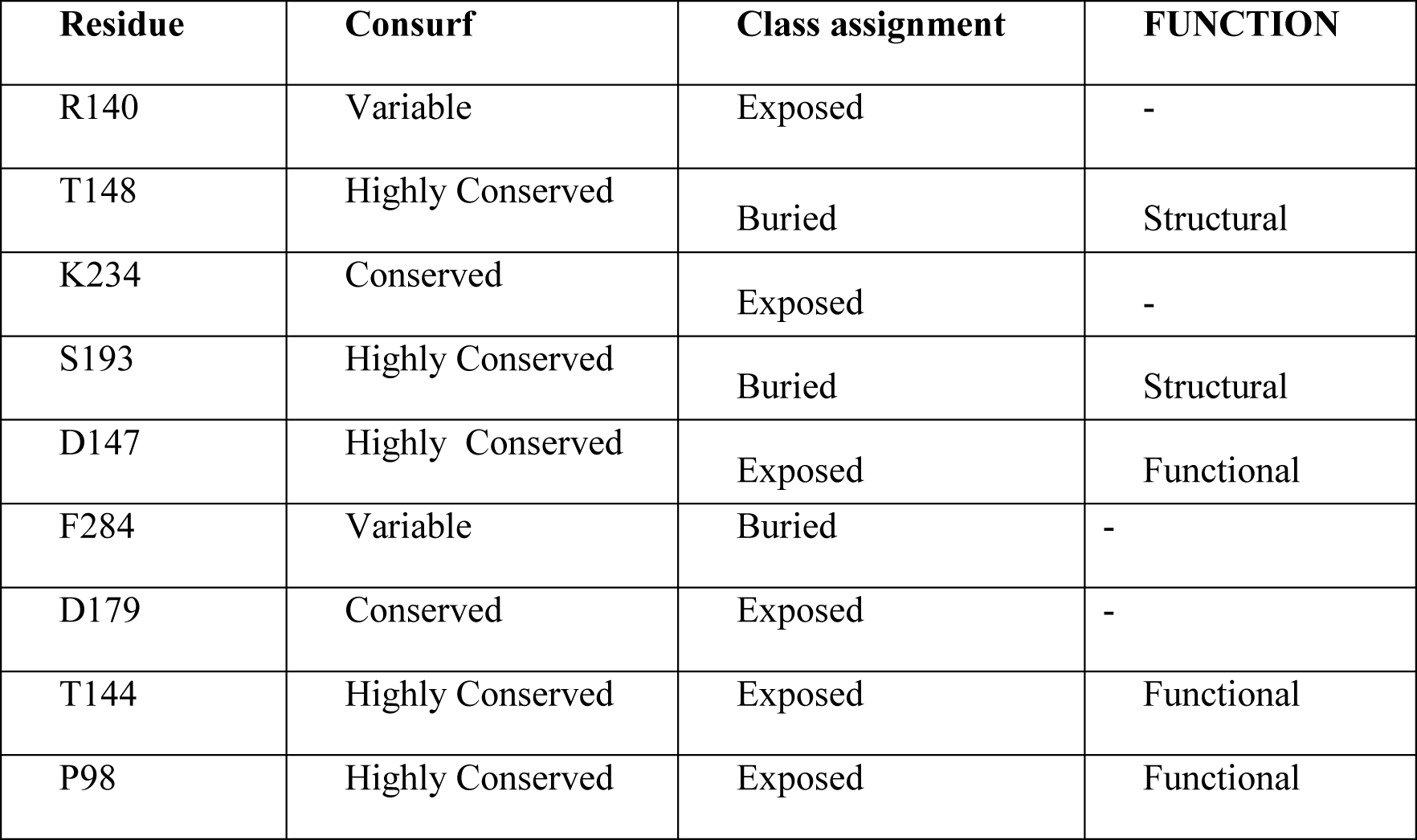
Conservation profile of amino acids in ADPRC 1.

| **Residue** | **Consurf** | **Class assignment** | **FUNCTION** |
| --- | --- | --- | --- |
| R140 | Variable | Exposed | - |
| T148 | Highly Conserved | Buried | Structural |
| K234 | Conserved | Exposed | - |
| S193 | Highly Conserved | Buried | Structural |
| D147 | Highly Conserved | Exposed | Functional |
| F284 | Variable | Buried | - |
| D179 | Conserved | Exposed | - |
| T144 | Highly Conserved | Exposed | Functional |
| P98 | Highly Conserved | Exposed | Functional |

#### Protein-protein interactions analysis

From STRING protein-protein interaction analysis, CD38 was predicted with the strong interactions with BST1, ENPP1, ENPP3, NADK, NAMPT, NMNAT1, NMNAT2, NMNAT3, NNMT and PNP.(Fig 2) The STRING interaction result was further validated by using KEGG pathways for CALR. Interestingly, same set of proteins were involved in metabolic pathways, including Nicotinate and nicotinamide metabolism, Riboflavin metabolism, Pyrimidine metabolism, Pantothenate and CoA biosynthesis, Purine metabolism, Starch and sucrose metabolism, Calcium signaling pathway, Oxytocin signaling pathway in addition to Salivary secretion and Pancreatic secretion. Our literature search demonstrated that Pancreatic secretion pathway (<u>hsa04972</u>) is involved in pathways of Type 2 diabetes mellitus. (Fig 3)

**Figure 2:**
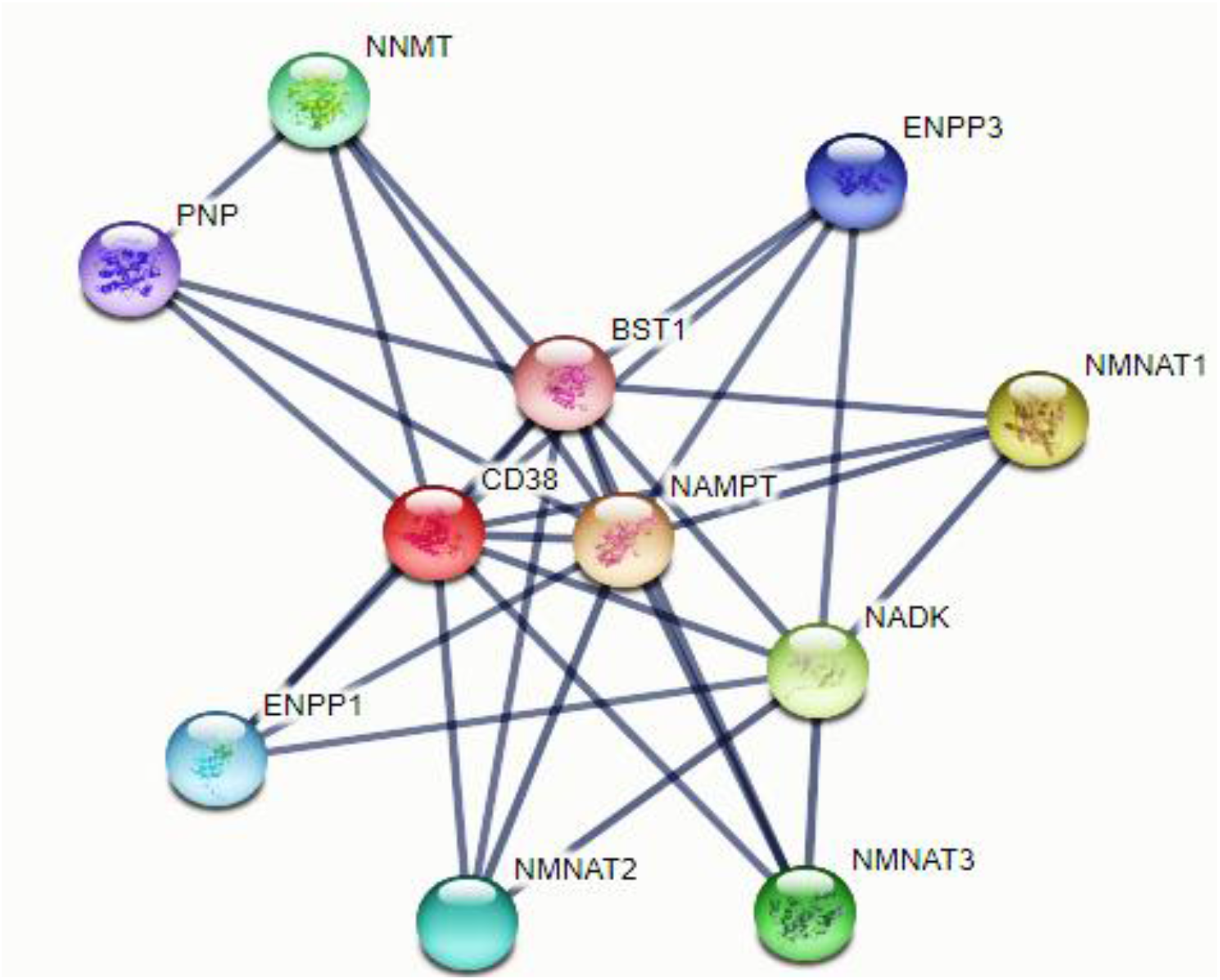
Interaction between CD38 and its related protein.

**Figure3:**
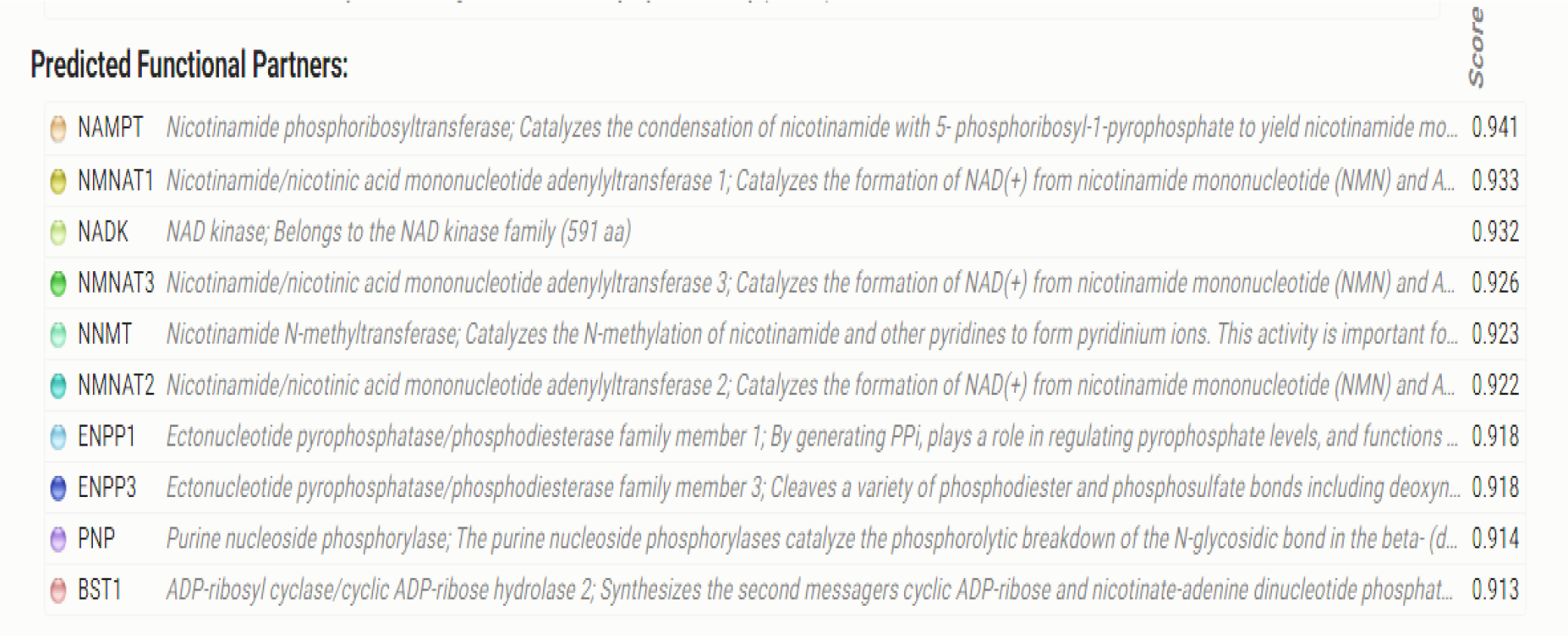
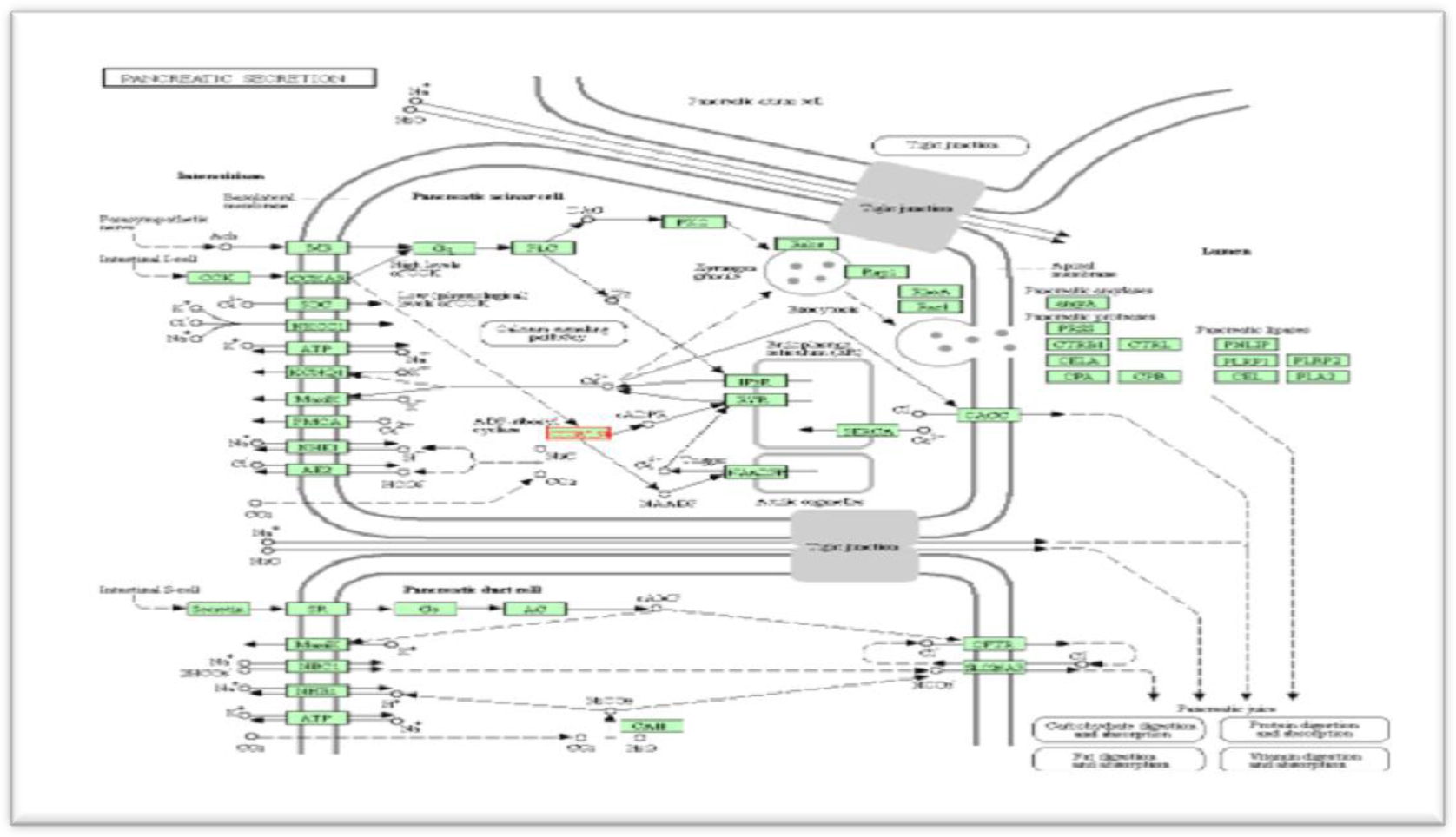
Pancreatic secretion pathway in Type 2 diabetes mellitus

##### Analysis of the effect of nsSNPs in ADPRC by MutPred server

The effect on the polymorphism in CD38 gene on ADPRC 1 protein was predicted using mutpred software which shows the following six SNPs (T148M, K234M, S193C, D147G, F284C, D179G, T144N, P998L) to be harmful in term of disturbing the molecular mechanisms, while the SNP R140W was predicted to be non-harmful with score of .275.(Table 6).

**Table 6:** Analysis of the effect of nsSNPs in ADPRC 1 structure, function, and evolution by MutPred server.

| Amino acid<br>Change | Score (g) | Prediction | Top 5 Molecular Molecular Mechanism Disrupted (P) |
| --- | --- | --- | --- |
| R140W | 0.275 | Non harmful | Loss of solvent accessibility (P = 0.086)<br>Loss of helix (P = 0.1706)<br>Gain of sheet (P = 0.1945)<br>Loss of catalytic residue at R140 (P = 0.2226)<br>Loss of disorder (P = 0.2286) |
| T148M | 0.778 | Highly harmful | Gain of catalytic residue at T148 (P = 0.0143)<br>Loss of sheet (P = 0.1158)<br>Gain of helix (P = 0.132)<br>Loss of phosphorylation at T144 (P = 0.614)<br>Loss of disorder (P = 0.6792) |
| K234N | 0.701 | Harmful | Loss of ubiquitination at K234 (P = 0.0116)<br>Loss of methylation at K234 (P = 0.0138)<br>Gain of loop (P = 0.0851)<br>Loss of disorder (P = 0.2145)<br>Loss of helix (P = 0.2662) |
| S193C | 0.900 | Highly harmful | Loss of MoRF binding (P = 0.0729)<br>Gain of methylation at K190 (P = 0.078)<br>Loss of ubiquitination at K190 (P = 0.1498)<br>Loss of phosphorylation at S193 (P = 0.219)<br>Loss of catalytic residue at S193 (P = 0.2472) |
| D147G | 0.863 | Highly harmful | Loss of stability (P = 0.1122)<br>Gain of catalytic residue at D147 (P = 0.1166)<br>Gain of sheet (P = 0.1945)<br>Loss of helix (P = 0.3949)<br>Loss of disorder (P = 0.6107) |
| F284C | 0.854 | Highly harmful | Gain of catalytic residue at L285 (P = 0.0249)<br>Gain of disorder (P = 0.0444)<br>Gain of ubiquitination at K289 (P = 0.0594)<br>Loss of MoRF binding (P = 0.1028)<br>Gain of sheet (P = 0.1208) |
| D179G | 0.687 | Harmful | Gain of MoRF binding (P = 0.065)<br>Gain of methylation at K178 (P = 0.0896)<br>Gain of relative solvent accessibility (P = 0.09)<br>Loss of disorder (P = 0.1701)<br>Loss of helix (P = 0.1706) |
| T144N | 0.785 | Highly harmful | Gain of relative solvent accessibility (P = 0.09)<br>Loss of sheet (P = 0.1158)<br>Gain of solvent accessibility (P = 0.1422)<br>Loss of catalytic residue at E146 (P = 0.2117)<br>Loss of loop (P = 0.2237) |
| P98L | 0.895 | Highly harmful | Gain of relative solvent accessibility (P = 0.0479)<br>Gain of solvent accessibility (P = 0.0704)<br>Loss of disorder (P = 0.0862)<br>Loss of methylation at K96 (P = 0.0868)<br>Gain of ubiquitination at K96 (P = 0.102) |

## Discussion

CD38 is known to be a direct contributor in pathogenesis of CLL but still the exact mechanism is poorly understood. CD38 expression was observed to increase in CLL. More ever CD38 consider as a dual function molecule, it can plays that role by acting as a receptor/adhesion molecule as well as an ecto-enzyme. Depending on the cell type as well as CD38 maturation state, the downstream signaling and biological consequences of CD38 activation will take place**^(40–42)^**. This can be attributed to the fact that CD38 often associates and acts in synergy with other cell type specific surface receptors. In CLL cells, CD38 collaborates with the BCR/CD19 complex for signal transduction. **^(43–44)^**

The present study revealed that CD38 gene polymorphisms (CD38Missense SNPs) may result in, harmful alteration on entire protein structure, stability and physiochemical properties. Subsequently, this change will initiate modification on multiple functions of molecule, in addition to interaction between CD38 and other molecules in wide variety pathways **^(45)^**. One Study, was performed by Fadl, et al **^(46)^** in Sudan and concluded that Missense CD38 SNP (rs1800561) (418C>T, Arg140Trp) has higher genotypic frequency than wild type allele (CC), among CLL Sudanese patients. On another hand, a significant correlation between CD38 SNP and age of patients was found in her work, with a higher frequent among patients whose their ages (≤60) years, Concerning gender of patients, they found that {SNP were higher in males than in females} even though, they were stated The following: CD38 gene polymorphism caused significant increase in some hematological parameters {TWBCS count(C/L), Absolute Lymphocytes and PLTS count(C/L)} but no significant correlation between the SNP and Hb conc(g/dl).TWBCS count(C/L) and Absolute Lymphocytes count(C/L) were strongly correlated with CD38 gene SNP, although it seems to be weakly correlated with PLTS count(C/L). Further study done by Jamroziak et al in Poland **^(47)^**. They described that frequency of wild type allele (CC) allele was higher than CD38 gene polymorphism (CT) with (P-value=0.97) in CLL patients, against low (rs1800561 CT allele) frequency, they did not detect any individuals homozygous for this allele, which is consistent with Fadl’s study. And it did not associate with age of patients, which is disagreed with Fadl’s findings. variation in sample size and study population might explain the gap of results between these studies.Other study in Poland, that was published by Szemraj-Rogucka et al **^(48)^**, stated that, The patients who were diagnosed with MM, Their median age never been differ significantly from wild-type(rs1800561) CC homozygotes(61 years) and others who have heterozygous (rs1800561) CT genotype (68 years)with P-value= 0.23.whereas, they were reported that no Significant association between CD38 genotype and age nor gender of MM patients. According to P-value = 0.74 for rs1800561 CT analysis. Additional study that investigated CD38 polymorphism as risk of diseases is a study which conducted in Spain by González-Escribano et al **^(49)^**.Their results did not support the hypothesis that the CD38 gene polymorphism is related to the risk of SLE, that suggests no individual who have (CT genotype) were detected among patients with SLE, nor the healthy individuals. These results, support the absence (or the extremely low frequency) of this polymorphism in Spanish population. Former study which carried out by Yagui et al **^(50)^** in Japan. Noticed that, there is an significant association between CD38 gene polymorphism and type II Diabetes mellitus. By way of initiating Considerable declining in both ADP-ribosyl cyclase and cADPR hydrolase activities by nearby 50%, that could contribute to the diminishing of (in vivo) insulin secretion in patients with Type II diabetes mellitus. It had been suggested that (CT) allele frequency were significantly different between the type II diabetes mellitus patients, as (P-value= 0.004). CD38 gene SNP may contribute to the development of type II diabetes mellitus in collaboration with other genetic defects in beta cell function and insulin action, this finding along support our Hypothesis, that CD38 gene polymorphisms could contributed to some diseases despite the fact that González-Escribano’s study was disagreed with it.(Figure3).

Twelve SNPs were found in this work to be functional deleterious by three soft wares, further analysis by the remnant software reveals nine of them were deleterious. Of these SNPs eleven, were novel(not reported before in the literature and one was reported in many studies as predisposing factor to some diseases. Likewise, stability analysis reported that Seven SNPs were decreased stability while two were increased stability. Even though, eight SNPs were decreased against one was increased by other software. Furthermore (rs 180056 1) SNP was reported before in the work of Fadl et al. where they found it higher in frequency than wild type allele that is suggested as risk factor for CLL. Project hope was used to predict the effect of these SNPs on the physiochemical prorates of the protein where it were found to effect the size, charge and hydrophobic state of the protein s=confirming the deleterious effect of these SNPs. Moreover the effect on molecular mechanism was predicted using MutPred server. Software which shows that all the SNPs were harmful on the molecular level except R140W which was un anticipated result as this SNPs was reported in CLL patients, which raise the need for more focus on this SNP on further research.Our findings are agree with Fadl’s while disagree with Jamroziak’s findings. Also, Consistent with Yagui’s and Inconsistent with Szemraj-Rogucka, González-Escribano regarding other diseases.It was reported in the literature that CD38 gene has complex relation with other genes. This relation was studied in this work by using STRING software, where it shows a complex interactions with other genes like PNP, BST1, NNMT, and many others.

## Conclusion

The bioinformatics analysis has got highest concern to screen diseases associated nsSNP at molecular level. In this study a comprehensive bioinformatics analysis has been performed to investigate the effect of nsSNPs on structure and function of CD38. As CD38 involved in many pathways, hence any changes on it structure and function would have an impact on many pathways, involved in wide verity diseases. Nine nsSNPs were predicted to be the most damaging mutations for CD38 leading to loss or disturbance of the protein internal and external interactions and eventually loss of the protein's function as well as disease association. This is computational approach, thus wet-lab based studies with clinical evidences are necessary to accompaniment the findings of this study.

## Recommendations

Additional wet-lab based researches are recommended to study outcomes that may occur as consequence of CD38 SNPs, which might amend other functions of CD38 molecule. As it was involved in many pathways that could end in wide verity diseases. Have a preference to use more soft wares, and to state advance study on a specific population, in purpose of taking genetic background of patients in consideration. As well as focusing on CD38 as flow marker and its impression on prognosis. Also, the effect of the CD38 genotype on the hematological findings has to be studied further, for the reason that those patients could be at risk of other haematopathological complications.

## Ethical Consideration

This study is a bioinformatic approach and doesnt required ethical approval.

## Author’s Contribution

This Study was fulfilled in collaboration among all authors. Author **H.A.O.F** designed the study, contributed in bioinformatic analysis, prepared and wrote the manuscript. Author **A.H.A.H** participated in the bioinformatic analysis, preparation and revision of manuscript. Author **S.G.E** established the protocol of the Study, accomplished the bioinformatic analysis and final revision of the study. All authors critically reviewed the manuscript and approved the final draft

## Acknowledgment

We would like to thank our families, frindes and teachers who always provid us a highly support.

## References

1. Xu W, Li JY, Miao KR, et al. The negative prognostic significance of positive direct antiglobulin test in Chinese patients with chronic lymphocytic leukemia. Leuk Lymphoma 2009;50:1482–7.

2. Keating MJ. Management of chronic lymphocytic leukemia: a changing field. Rev Clin Exp Hematol 2002;4:350–65. In addititon of Diagnisic and prog…risk factor or predisposing factor.

3. Rozman C, Montserrat E. Chronic lymphocytic leukemia. N Engl J Med 1995;1333:1052–57.

4. Zwiebel JA, Cheson BD. Chronic lymphocytic leukemia: staging and prognostic factors. Semin Oncol 1998;25:42–59.

5. Mauro FR, Foa R, Giannarelli D, et al. Clinical characteristics and outcome of young chronic lymphocytic leukemia patients: a single institution study of 204 cases. Blood 1999;94:448–54.

6. Zeeshan R, Sultan S, Irfan SM, Kakar J, Hameed MA. Clinico-hematological profile of patients with B-chronic lymphoid leukemia in Pakistan. Asian Pac J Cancer Prev 2015;16:793–6. And. Rossi D, Gaidano G, The clinical implications of gene mutations in chronic lymphocytic leukaemia. Br J Cancer. 2016 Apr 12;114(8):849–54.2016.78. Epub 2016 Mar 31.

7. Houlston RS, Catovsky D, Yuille MR. Genetic susceptibility to chronic lymphocytic leukemia. Leukemia 2002;16:1008–14.

8. Zucchetto A, Vaisitti T, Benedetti D, Tissino E, Bertagnolo V, et al. (2012) The CD49d/CD29 complex is physically and functionally associated with CD38 in B-cell chronic lymphocyt ic leukemia cells. Leukemia 26: 1301–1312.

9. Ferrero E, Saccucci F, Malavasi F. The human CD38 gene: polymorphism, CpG island, and linkage to the CD157 (BST-1) gene. Immunogenetics 1999;49:597–604.

10. Crystal Structure of Human CD38 Extracellular Domain Liu. Q,Kriksunov. I.A, Graeff. R, Munshi. C, Lee. H.C, and Hao. Q,Structure, Vol. 13, 1331–1339, September, 2005, DOI 10.1016/j.str.2005.05.012

11. Deaglio S, Vaisitti T Atlas Genet Cytogenet Oncol Haematol. 2012; 16(7) 448.

12. Damle RN, Wasil T, Fais F, Ghiotto F, Valetto A, Allen SL, Buchbinder A, Budman D, Dittmar K, Kolitz J, Lichtman SM, Schulman P, Vinciguerra VP, Rai KR, Ferrarini M, Chiorazzi N. Ig V gene mutation status and CD38 expression as novel prognostic indicators in chronic lymphocytic leukemia. Blood, 94(6), 1999, 1840–1847.

13. Deaglio S, Vaisitti T, Aydin S, Ferrero E, Malavasi F. In-tandem insight from basic science

14. Deaglio S, Vaisitti T, Billington R, Bergui L, Omede P, et al. (2007)CD38/CD19: a lipid raft-dependent signaling complex in human Bcells. Blood 109: 5390–5938.

15. Susceptibility genes and B-chronic lymphocytic leukaemia Susan L Slager et al. Br J Haematol. 2007 Dec.

16. Deaglio. S. et al. Blood. 2006.In-tandem insight from basic science combined with clinical research: CD38 as both marker and key component of the pathogenetic network underlying chronic lymphocytic leukemia

17. Abramenko. I.V et al. Leuk Res. 2012 Oct.CD38 gene polymorphism and risk of chronic lymphocytic leukemia.

18. Genetic Variants of Snps RS6449182 CD38 Gene: Correlation with Age, Sex, and Tumor Stage in Patients with Colorectal Cancer. December OM&P 2016Volume 2 Issue 3, 4 omp2016.003.0036 http://www.operamedphys.org/OMP_2016_03_0036

19. Jamroziak. K et al CD38 gene polymorphisms contribute to genetic susceptibility to B-cell chronic lymphocytic leukemia: evidence from two case-control studies in Polish Caucasians. Cancer Epidemiol Biomarkers Prev. 2009 Mar. 18(3):945–53.

20. “single-nucleotide polymorphism / SNP | Learn Science at Scitable”. www.nature.com. Archived from the original on 2015-11-10. Retrieved 2015-11-13.

21. Cancer Epidemiol Biomarkers Prev 2009;18(3):945–53)combined with clinical research: CD38 as both marker and key component of the pathogenetic network underlying chronic lymphocytic leukemia. Blood 2006;108:1135–44.

22. Ferrero. E, Saccucci. F, Malavasi. F. The human CD38 gene:polymorphism, CpG island, and linkage to the CD157 (BST-1) gene.Immunogenetics 1999;49:597–604.

23. Wang. S, Zhu. W, Wang. X, Li J, Zhang K, et al. (2014) Design, synthesis and SAR studies of NAD analogues as potent inhibitors towards CD38 NADase. Molecules 19: 15754–15767.

24. Moreau C, Liu Q, Graeff R, Wagner GK, Thomas MP, et al. (2013) CD38 Structure-Based Inhibitor Design Using the 1-Cyclic Inosine 5’-Diphosphate Ribose Template. PLoS One 8:e66247.

25. Kellenberger E, Kuhn I, Schuber F, Muller-Steffner H (2011) Flavonoids as inhibitors of human CD38. Bioorg Med Chem Lett 21: 3939–3942.

26. Zhou Y, Ting KY, Lam CM, Kwong AK, Xia J, et al. (2012) Design, synthesis and biological evaluation of noncovalent inhibitors of human CD38 NADase. Chem Med Chem 7: 223–228.

27. Wall KA, Klis M, Kornet J, Coyle D, Ame JC, et al. (1998) Inhibition of the intrinsic NAD+ glycohydrolase activity of CD38 by carbocyclic NAD analogues. Biochem J 335: 631–636.

28. Swarbrick JM, Graeff R, Zhang H, Thomas MP, Hao Q, et al. (2014) Cyclic adenosine 5’-diphosphate ribose analogs without a “southern” ribose inhibit ADP-ribosyl cyclase-hydrolase CD38. J Med Chem 57:8517–8529.

29. Sauve AA, Schramm VL (2002) Mechanism-based inhibitors of CD38:a mammalian cyclic ADP-ribose synthetase. Biochemistry 41: 8455–8463.

30. Becherer JD, Boros EE, Carpenter TY, Cowan DJ, Deaton DN, et al.(2015) Discovery of 4-Amino-8-quinoline Carboxamides as Novel, Submicromolar Inhibitors of NAD-Hydrolyzing Enzyme CD38. J MedChem 58: 7021–7056.

31. Kwong AK, Chen Z, Zhang H, Leung FP, Lam CM, et al. (2012)Catalysis-based inhibitors of the calcium signaling function of CD38.Biochemistry 51: 555–564.

32. Kirchberger T, Wagner G, Xu J, Cordiglieri C, Wang P, et al.(2006) Cellular effects and metabolic stability of N1-cyclic inosine diphosphoribose and its derivatives. Br J Pharmacol 149: 337–344.

29. Adzhubei I, Jordan DM, Sunyaev SR. Predicting functional effect of human missense mutations using PolyPhen-2. Curr Protoc Hum Genet. 2013;Chapter 7:Unit7 20.

30. Ng PC, Henikoff SJGr. Accounting for human polymorphisms predicted to affect protein function. 2002;12(3):436–46.

31. Manickam M, Ravanan P, Singh P, Talwar P. In silico identification of genetic variants in glucocerebrosidase (GBA) gene involved in Gaucher's disease using multiple software tools. Front Genet. 2014;5:148.

32. Ng PC, Henikoff SJARGHG. Predicting the effects of amino acid substitutions on protein function. 2006;7:61–80.

33. Schneider G, Hu J, Sim N-L, Kumar P, Henikoff S, Ng PC.

34. Choi Y, Sims GE, Murphy S, Miller JR, Chan AP. Predicting the functional effect of amino acid substitutions and indels. PLoS One. 2012;7(10):e46688.

35. Kumar P, Mahalingam K. In silico approach to identify non-synonymous SNPs with highest predicted deleterious effect on protein function in human obesity related gene, neuronal growth regulator 1 (NEGR1). 3 Biotech. 2018;8(11):466.

36. Ramensky V, Bork P, Sunyaev S. Human non-synonymous SNPs: server and survey. Nucleic acids research. 2002 Sep 1;30(17):3894–900.

37. Adzhubei IA, Schmidt S, Peshkin L, Ramensky VE, Gerasimova A, Bork P, et al. A method and server for predicting damaging missense mutations. Nature methods. 2010;7(4):248–9.

38. Calabrese R, Capriotti E, Fariselli P, Martelli PL, Casadio RJHm. Functional annotations improve the predictive score of human disease-related mutations in proteins. 2009;30(8):1237–44.

39. López-Ferrando V, Gazzo A, de la Cruz X, Orozco M, Gelpí JLJNar. PMut: a web-based tool for the annotation of pathological variants on proteins, 2017 update. 2017;45(W1):W222–W8.

40. Capriotti E, Fariselli P, Casadio RJNar. I-Mutant2. 0: predicting stability changes upon mutation from the protein sequence or structure. 2005;33(suppl_2):W306–W10.

41. Venselaar H, te Beek TA, Kuipers RK, Hekkelman ML, Vriend GJBb. Protein structure analysis of mutations causing inheritable diseases. An e-Science approach with life scientist friendly interfaces. 2010;11(1):548.

42. Warde-Farley D, Donaldson SL, Comes O, Zuberi K, Badrawi R, Chao P, et al. The GeneMANIA prediction server: biological network integration for gene prioritization and predicting gene function. 2010;38(suppl_2):W214–W20.

43. Wang S, Li W, Liu S, Xu J. RaptorX-Property: a web server for protein structure property prediction. Nucleic acids research. 2016;44(W1):W430–5.

44. Pettersen EF, Goddard TD, Huang CC, Couch GS, Greenblatt DM, Meng EC, et al. UCSF Chimera—a visualization system for exploratory research and analysis. 2004;25(13):1605–12.

45. Gonzalez-Perez A, Lopez-Bigas N. Improving the assessment of the outcome of nonsynonymous SNVs with a consensus deleteriousness score, Condel. (1537-6605 (Electronic)).

46. Fadl. H and Humeida. A: Study of CD38 Gene Polymorphism in Sudanese Patients with CLL, Academic Journal of Cancer Research 13 (1): 08–13, 2020, DOI: 10.5829/idosi.ajcr.2020.08.13.

47. Jamroziak. K, Z. Szemraj, O.G. Izydorczyk, J. Szemraj, M. Bieniasz, B. Cebula,2009. Krzyszt of Giannopoulos, Ewa Balcerczak, Dorota Jesionek-Kupnicka, Malgorzata Kowal, Aleksandra Kostyra, Malgorzata Calbecka, Ewa Wawrzyniak, Marek Mirowski, Radzislaw Kordek, and Tadeusz Robak,2009. CD38 Gene Polymorphisms Contribute to Genetic Susceptibility to B-Cell Chronic Lymphocytic Leukemia: Evidence from Two Case-Control Studies in Polish Caucasians/ Cancer Epidemiology, Biomarkers & Prevention,March.Volume 18, Issue 3.

48. Zofia Szemraj-Rogucka Janusz Szemraj, Olga Grzybowska-Izydorczyk,Tadeusz Robak, Krzysztof Jamroziak: CD38 gene polymorphisms and genetic predisposition to multiple myeloma, acta haematologica poloncia 44 (2013) 58–62.

49. González-Escribano. M, F. Aguilar, B. Torres,J. Sánchez-Román and A. Núñez-Roldán,2004.CD38 polymorphisms in Spanish patients with systemic lupus erythematosus Author links open overlay pane, 2004.Servic iode Inmunología, Sevilla, Spain Unidad de Colagenosis, HU Virgen del Rocío, Servicio Andaluz de Salud, Sevilla and Spain,2004.

50. Yagui. K, F. Shimada, M. Mimura, N. Hashimoto, Y. Suzuki, Y. Tokuyama, K. Nata, A. Tohgo, F. Ikehata, S. Takasawa, H. Okamoto, H. Makino, Y. Saito and A. Kanatsuka,1998. A missense mutation in the CD38 gene, a novel factor for insulin secretion: association with Type II diabetes mellitus in Japanese subjects and evidence of abnormal function when expressed in vitro, Diabetologia 41: 1024±1028.

